# Mental travel in the social domain

**DOI:** 10.1101/849497

**Authors:** Mordechai Hayman, Shahar Arzy

## Abstract

“Mental travel” is the ability to imagine oneself in different places and times and to adopt other people’s point of view (POV), also termed “Theory of Mind (ToM)”. While ToM has been extensively investigated, self-projection with respect to ones’ own and others’ social networks has yet to be systematically studied.

Here we asked participants to “project” themselves to four different POVs: a significant other, a non-significant other, a famous-person, and their own-self. From each POV they were asked to rate the level of affiliation (closeness) to different individuals in the respective social network while undergoing functional MRI.

Participants were always faster making judgments from their own POV compared to other POVs (self-projection effect) and for people who were personally closer to their adopted POV (self-reference effect). Brain activity at the medial prefrontal and anterior cingulate cortex in the self POV condition was found to be higher compared to all other conditions. Activity at the right temporoparietal junction and medial parietal cortex was found to distinguish between the personally related (self, significant- and non-significant others) and unrelated (famous-person) individuals within the social network. Regardless of the POV, the precuneus, anterior cingulate cortex, prefrontal cortex, and temporoparietal junction distinguished between relatively closer and distant people. Representational similarity analysis (RSA) implicated the left retrosplenial cortex as crucial for social distance processing across all POVs.

## Introduction

“Mental time travel” (MTT) refers to the human capacity to intentionally disengage from the present moment and mentally wander to a specific past event or an event which may occur in the future (Tulving, 1985). One approach to the research of this well-studied cognitive operation, viewed MTT as a mechanistic “projection” of oneself on a so-called “mental-time-line” (Arzy et al., 2008, 2009; Buckner & Carroll, 2007). Furthermore, this neurocognitive mechanism was suggested to encompass not only the ability to shift one’s self-location in mental time but also in mental space, an ability crucial to imagining oneself in the place where an event has happened (Buckner & Carroll, 2007; Gauthier & van Wassenhove, 2016b; Tulving, 1985). It has been speculated that in the same way a person may “project” herself to another time and location, she may also shift her point-of-view (POV) to that of another person. Such a “projection” in the social domain is closely related to the concept of Theory of Mind (ToM) – the ability to consider others’ perspectives and mental states (Frith & Frith, 2003; Leslie, 1987; Premack & Woodruff, 1978; Buckner & Carroll, 2007).

The hypothesis that self-projection in the domains of space, time and person rely on similar neurocognitive systems received support, as they were found to share overlapping and adjacent brain activations, mainly in the medial parietal cortex, temporo-parietal junction and medial prefrontal cortex (Buckner & Carroll, 2007; Carrington & Bailey, 2009; Gauthier, Pestke, & van Wassenhove, 2018; Gauthier & van Wassenhove, 2016b). Notably, these regions are closely related to the default network (DN) that supports processing of internally constructed representations (Buckner et al, 2008; Buckner & DiNicola, 2019). Remarkably, recent evidence points on the existence of parallel but distinct sub-networks in the DN, in particular separate sub-networks for processing information about people, episodes and places (Buckner & DiNicola, 2019; DiNicola, Braga, & Buckner, 2019; Silson et al., 2019).

In both the temporal and spatial domain, two main effects have been revealed: the first – self-projection – involves the imagination of oneself in a place or time beyond the here and now (Arzy et al., 2009, 2008; Buckner & Carroll, 2007; Gauthier & van Wassenhove, 2016a, 2016b), the second – self-reference – involves the determination of the temporal or spatial relations between different events and locations (respectively) to the “projected” self (e.g. whether specific life events have happened before or after this specific time (Anelli, Ciaramelli, Arzy, & Frassinetti, 2016a; Arzy et al., 2009, 2008; for self-reference in memory see Klein & Kihlstrom, 1986; Symons & Johnson, 1997) or whether a specific place is located east or west to the imagined location (Gauthier et al., 2018; Gauthier & van Wassenhove, 2016a, 2016b). These effects have not yet been directly investigated in the social domain. Recent research which has focused on the network character of this domain (the so-called “social network”) (Parkinson, Kleinbaum, & Wheatley, 2017; Weaverdyck & Parkinson, 2018) as well as a potential metric basis of this domain (Schafer & Schiller, 2018; Tavares et al., 2015) supports taking such an approach. To investigate self-projection in the person domain we used an ecologically-valid paradigm and functional MRI. We examined self-projection to POVs of other persons varying in levels of proximity to the experiencer (self, significant-other, non-significant other, famous-person) (Laurita, Hazan, & Spreng, 2018; Wlodarski & Dunbar, 2016), as well as self-reference to different people in these persons’ social network.

## Methods

Subjects. Fifteen healthy participants (nine females, mean-age: 25.5±3.1 years old) took part in the study. All participants had normal or corrected to normal vision. Participants with a history of neurological or psychiatric treatment as well as those using psychoactive medications were excluded. All participants provided written informed consent, and the study was approved by the ethical committee of the Hadassah Medical Center.

Stimuli and procedure. At least a week prior to the experiment, participants were required to report the social relations of themselves, as well as those of three well-acquainted individuals varying in levels of proximity: a significant other (e.g. parent, spouse), a non-significant other (e.g. class-mate) and a famous-person (Israeli PM Benjamin Netanyahu). Namely, participants were asked to “project” themselves to each of these individuals own point of view (POV), generate a list of people that this individual is familiar with and rate the social distance from each person on the list on a 1-to-6 scale. This gave rise to a set of 16 names (“social network”) for each POV, divided in between 6 levels of personal proximity.

In the experimental task, participants were first presented with a target stimulus indicating the person whose POV they should adopt (e.g. “Netanyahu”), followed by pairs of stimuli consisting names of people from this person’s social network (Fig. S1A). For each pair of names, participants were instructed to determine which of the two is personally closer to “themselves” (Peer et al., 2015, 2019), according to their current POV. Responses were recorded by pressing the corresponding (left or right) buttons. Stimuli were presented in a randomized block design. Each block started by presentation of a target stimulus for 5 seconds, followed by consecutive presentation of four stimuli pairs, each for 2.5 s (Fig. S1B). Each block (10s) was followed by 5s of fixation. Participants were instructed to respond accurately but as fast as possible. A 4-min training task was delivered before the experiment. The experiment consisted of six experimental runs for each participant, each run containing 8 blocks in a randomized order (two blocks for each POV). Stimuli were presented using the Presentation® software (Version 18.3, Neurobehavioral Systems, Inc., Berkeley, CA, www.neurobs.com).

Analysis of behavioral data. Stimuli pairs were categorized in 3 subgroups according to their relative distance – close, distant and mixed: “close” - indicating both stimuli were rated 1-3 from the relevant POV; “distant” - indicating both stimuli were rated 4-6; “mixed” indicating one stimulus was rated 1-3 and the other 4-6. Stimuli pairs did not include adjacent levels of proximity and were equally distributed between levels of proximity. Linear mixed-effects (LME) models were fitted for the participants’ response times (RTs). As fixed factors in the LME, we tested effects of self-projection (POV; self, significant other, nonsignificant other and famous-person), self-reference (distance; close, distant, mixed) and disparity (disparity; the difference between the closeness ratings of the stimuli pair; 2-5). The model was controlled for within-subject effects by including random effects for participants (participants; 15 levels). To identify the model that best describes our results and avoid overfitting, we used a statistical model selection strategy based on Akaike information criterion (AIC) score (Akaike, 1998). The model with the lowest AIC was chosen (likelihood-ratio test; p<0.05; Table 1). Bonferroni corrected post hoc tests were then derived from the selected final model between all pairs. Behavioral data analysis was performed using the SPSS software (IBM SPSS Statistics, Version 22.0. Armonk, NY: IBM Corp.).

**Table 1.**
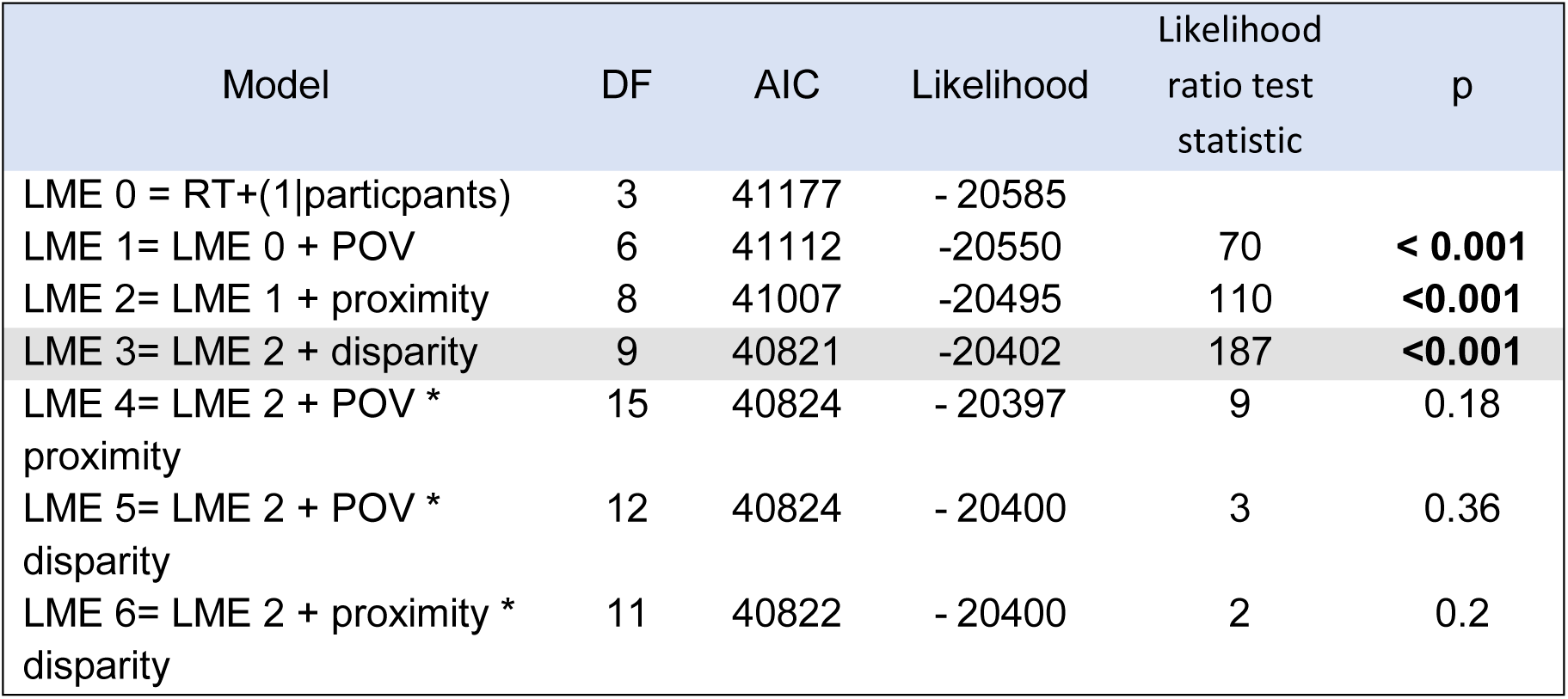
Linear mixed-estimates (LME) models. The different potential models are shown, as based on permutations of performance (response times, RTs), the adopted POV, the different levels of proximity and the disparity between the people that were presented in the stimuli. Models showing significant effects, as compared to a previous iteration, were chosen (p-values are provided for the likelihood ratio test). LME 4 is the model used in our analysis (DF = degrees of freedom; AIC = Akaike information criterion).

MRI acquisition. Subjects were scanned in a 3T Siemens Skyra MRI (Siemens, Erlangen, Germany) at the Edmund and Lily Safra Center (ELSC) neuroimaging unit. Blood oxygenation level-dependent (BOLD) contrast was obtained with an echo-planar imaging sequence [repetition time (TR), 2500 ms; echo time (TE), 30 ms; flip angle, 75°; field of view, 192 mm; matrix size, 64 × 64; functional voxel size, 3 × 3 × 3 mm; 44 slices, descending acquisition order, no gap]. In addition, T1-weighted high resolution (1 × 1 × 1 mm, 160 slices) anatomical images were acquired for each subject using the MPRAGE protocol [TR, 2500 ms; TE, 2.98 ms; flip angle, 9°; field of view, 256 mm].

MRI preprocessing. fMRI data were analyzed using the SPM 12 software package (Friston, Ashburner, Kiebel, Nichols, & Penny, 2007) and in-house Matlab (Mathworks) scripts (available at neuropsychiatrylab.com). Preprocessing of functional scans included slice timing correction (sinc interpolation), 3D motion correction by realignment to the first run image (2nd degree B-Spline interpolation), exclusion of runs with maximal motion above a single voxel size (3 mm) in any direction, smoothing (full width at half maximum (FWHM) = 4 mm), and co-registration to the anatomical T1 images. Anatomical brain images were corrected for signal inhomogeneity and skull-stripped. All images were subsequently normalized to Montreal Neurological Institute (MNI) space (3 × 3 × 3 mm functional resolution, 4th degree B-Spline interpolation).

## Functional MRI analyses

### Parameter estimation analysis

To assess the selective activations elicited by different experimental conditions (self-projection to the different POVs and self-reference to relatively “close” and “distant” people), we applied a general linear model (GLM) analysis. Predictors were constructed for all conditions, convoluted with a canonical hemodynamic response function, and the model was independently fitted to the time course of each voxel. Motion parameters with their derivatives and the corresponding squared items were added to the GLM to remove motion-related noise (Friston et al., 1996). For each participant, we averaged the GLM estimated parameter value (‘beta’ values) for each condition across each regions of interest (ROI, see below). Repeated measure ANOVA were run between conditions across all participants (p < 0.05, FDR-corrected).

### MVPA Searchlight Analysis

To localize brain regions with sensitivity to the task’s conditions (self-projection and self-reference), we performed a searchlight analysis (Kriegeskorte, Goebel, & Bandettini, 2006; Z-score = 2.58 (p<0.005)). In this analysis, classification is performed with a support vector machine (SVM) classification algorithm implemented in the CoSMoMVPA toolbox for Matlab (Maris & Oostenveld, 2007; Oosterhof, Connolly, & Haxby, 2016). The searchlight analysis was performed on the GLM beta values, across MNI-normalized whole brain. Each voxel was taken as a center point for a sphere with a radius of 3 voxels (up to 123 voxels in the Searchlight sphere). For each of those spheres, a 5-fold leave-one-out cross-validation was performed, resulting in 1 accuracy value per sphere. The classification accuracy of a given voxel was calculated as the average accuracy of all spheres that included this voxel. To assess the statistical significance of searchlight maps across participants, all maps were corrected for multiple comparisons without choosing an arbitrary uncorrected threshold using threshold-free cluster enhancement (TFCE) on the cluster level (Smith & Nichols, 2009). A Monte-Carlo simulation permuting condition labels was used to estimate a null TFCE distribution. First, 100 null searchlight maps were generated for each participant by randomly permuting condition labels within each obtained searchlight classification. Next, 100,000 null TFCE maps were constructed by randomly sampling from these null data sets in order to estimate a null TFCE distribution (Stelzer, Chen, & Turner, 2013), obtaining a group level Z-score map of the classifier results.

To examine the structure of the classification errors, we have run an SVM classifier for each ROI over all voxels and calculated a confusion matrix (CM) of the MVPA classifier. The classifier cross-correlation method produced six predicted labels for each condition pattern. To determine the extent of each condition in each ROI, we computed F1-scores for the classification results (Chinchor & Nancy, 1992; Sokolova, Japkowicz, & Szpakowicz, 2006). Repeated measure ANOVA were run between the F1 scores for all conditions as well as a null score of 0.25, over all participants (p < 0.05, FDR-corrected). Percentage of the classifier’s guessed trials were averaged across all participants.

### Representational similarity analysis (RSA)

A representational similarity analysis (Kriegeskorte et al., 2008) was used to “probe” brain regions whose similarity structure of local BOLD response reflected each participant’s social distance ratings from each tested POV. At each searchlight center, a nonparametric test of representational relatedness was performed on beta values from GLM analysis to evaluate the significance of the correlation between behavioral and neural representational dissimilarity matrices (RDMs). This procedure allows for each participant’s cortex to be continuously mapped in terms of the relatedness of representations manifested in local fMRI responses and behavioral ratings. This approach therefore provides a direct assessment of the level in which each participant’s BOLD signal within each searchlight sphere capture the information contained the social distances between POVs as reported by participants. This procedure produces a map with a scalar correlation value between the behavioral and neural RDMs for each participant. Group level analysis has been run in the same manner as for the MVPA analysis.

## Results

Behavioral results. Post hoc analysis on the linear mixed model estimates showed response times (RT) for judgements made from the participants’ own POV (1302 ± 339ms; mean ± SD) as well as the significant others’ POV (1357 ± 324ms) were significantly faster than those made from the non-significant others’ POV (1439 ± 359ms) and the famous-person POV (1432 ± 357ms; F=17.8, p<0.001, Sidak corrected; Fig. 1A). These findings suggest self-projecting to a personally distant individuals’ POV bears a cognitive cost reflected in an extended RT, which is less apparent when self-projecting to a personally close individual. Moreover, for all four POV conditions, judging between people who were personally close (1287 ± 338, 1294 ± 302, 1410 ± 374, 1399 ± 391 ms, respectively) resulted in significantly faster RTs than judging between people who were personally distant (1473 ± 352, 1506 ± 313, 1547 ± 336, 1548 ± 323 ms, respectively; F=59.3, p < 0.001, Sidak corrected; Fig. 1B), implying a self-reference effect.

**Figure 1.**
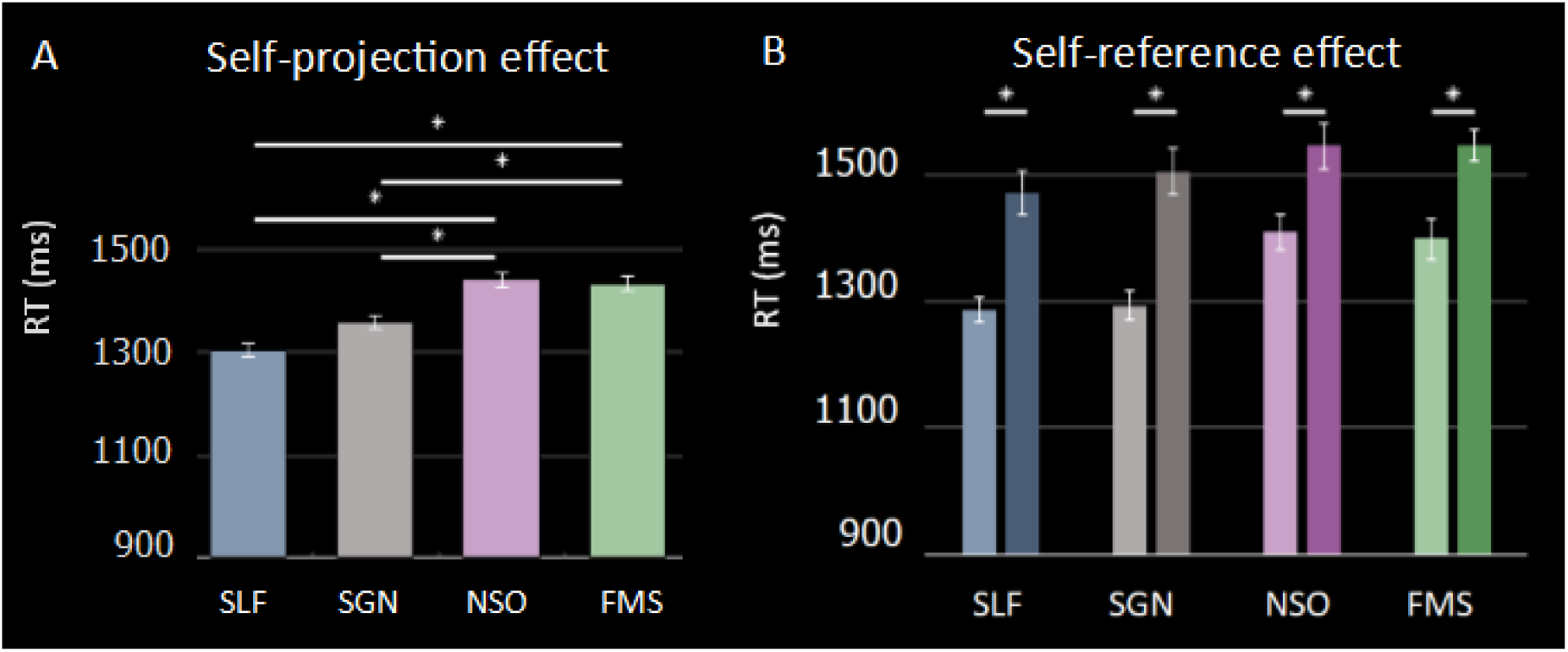
Behavioral results. (A) Self projection effect. Response times (RTs) are plotted separately for the different points of view (POV) conditions: self (blue), significant other (grey), non-significant other (purple) and famous-person (green). Note the significantly higher RTs for, non-significant other and famous-person, with respect to the self and significant other. (B) Self reference effect. For all POV conditions RTs were higher for the relatively distant stimuli pairs (right bars, darker colors) than the relatively closer ones (left bars, lighter colors). Error bars indicate standard error of the mean (SEM).

## Neuroimaging results

### ROI identification

Group analysis on the searchlight MVPA results (spheres sensitive for POV) revealed six functional clusters containing information regarding the adopted POV (Fig 2A, defined as ROIs for further analysis, according to the AAL2 atlas (Rolls, Joliot, & Tzourio-Mazoyer, 2015; Tzourio-Mazoyer et al., 2002)): (1) the medial parietal cortex – including the precuneus bilaterally and the left posterior cingulate cortex (PCC); (2) the right temporo-parietal junction (TPJ) including the right angular cortex and the middle temporal gyrus; (3) the left TPJ including the supramarginal, angular and superior temporal gyri; (4) the left inferior frontal lobe (IFL); (5) the right superior frontal lobe (SFL); and (6) a medial frontal cortex (mFC) including the medial prefrontal and the anterior cingulate cortices (ACC) bilaterally and part of the left SFL.

**Figure 2.**
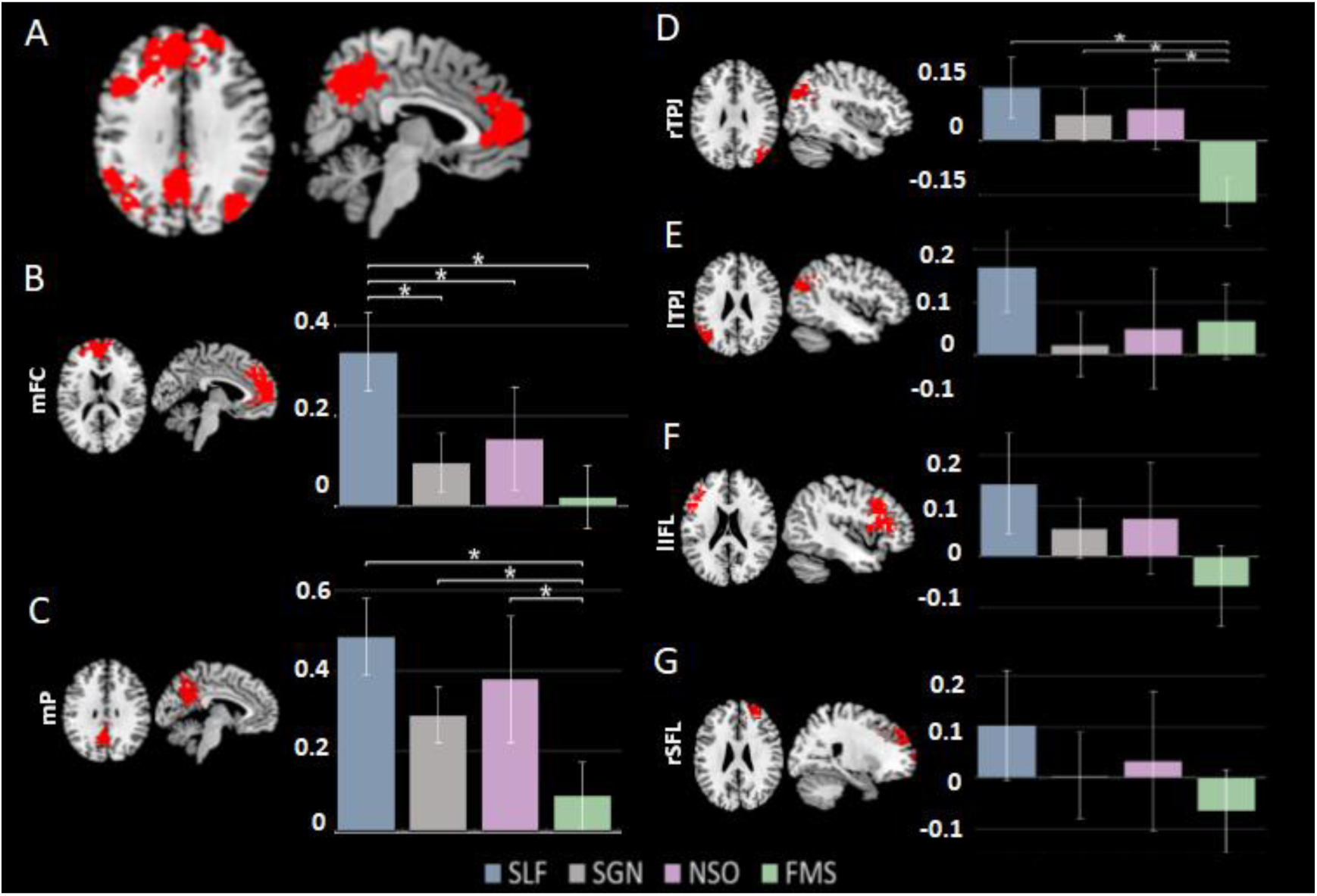
Neuroimaging results. (A) Axial (left) and mid-sagittal (right) images depicting results of searchlight-based multi-voxel pattern analysis (MVPA), classifying between the four POV conditions (p<0.005, TFCE corrected). (B-G). The six ROIs used in the parameter estimates analysis are shown (left). GLM parameters for each ROI are shown (right) for each POV separately, averaged across subjects. Significant differences are indicated by asterisks (p<0.05 FDR corrected; mFC – medial frontal cortex, mP – medial parietal, TPJ – temporoparietal junction, IFL – inferior frontal lobe, SFL – superior frontal lobe, r – right, l- left).

Searching for spheres sensitive to stimuli distance (close vs. distant stimuli) revealed activations at the medial prefrontal cortex, anterior and middle cingulate cortex, precuneus, cuneus, the TPJ and mFC bilaterally; and the right supramarginal gyrus, reflecting the self-reference effect (Fig. S2).

### Parameter estimation analysis

Parameter estimation analysis was used to specify values of activation in each ROI. The medial frontal ROI (Cluster 6) showed higher activity for the self POV condition than all other POV conditions (p < 0.05 for all comparisons, FWER corrected; Fig. 2B). Brain activation at the medial parietal and the right TPJ ROIs (Clusters 1, 2) distinguished between the POV conditions that are personally closer to self (self, significant other and non-significant other) and the POV condition that is personally distant (famous-person) (Fig. 2C,D). The left TPJ, left IFL and right SFL ROIs (Clusters 3, 4, 5) were non-significant for this distinction (Fig. 2E,F,G). These results were also corroborated in a confusion matrix analysis of the different ROIs (Table S2). For all ROIs, the activity pattern for the famous-person POV condition was significantly distinguishable from all other POV conditions. While the self POV condition showed a significant distinguishable in the left IFL, right SFL, right TPJ and medial parietal clusters, the significant other POV condition showed significant distinguishable in the left and right TPJ, and the non-significant other POV condition did not show any significant distinguishable (Table S2, Figure S2). Finally, application of RSA on the searchlight results in the group level identified one cluster (35 voxels, p<0.05) in the inferior part of the left retrosplenial cortex (Brodmann area 30, peak voxel at MNI coordinates −9,−55,10) in which activation patterns for the POV conditions significantly correlated with the social distances (Fig. 3).

**Figure 3.**
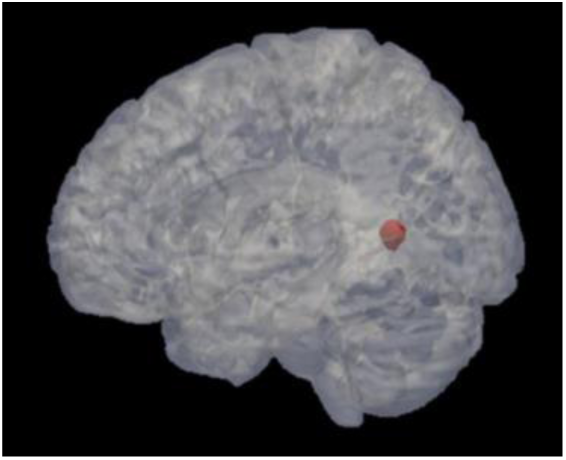
Representational similarity analysis (RSA) yielded a brain region within the retrosplenial cortex correlated with social distance. Searchlight method was used to investigate spheres around every voxel in the brain. At each sphere, a neural representational dissimilarity matrices (RDM) was generated based on pairwise correlation distances between local neural response patterns to each condition. Behavioral RDMs were constructed using Euclidean distances, based on the social distances between POVs, as reported by the participants, and correlated with local neural RDMs at each sphere. 35 voxels in the left precuneus (in red) showed significant correlation between the RDMs (p<0.05, TFCE correction with 10,000 iterations, shown on a 3D ‘glass brain’).

## Discussion

Behavioral analysis revealed adopting the point of view (POV) of others is facilitated when adopting POVs of personally closer individuals (self-projection effect). Furthermore, judging the social distance of people from different POVs was facilitated when judging between two personally close people relative to the adopted POV compared to judgment between two personally distant people (self-reference effect). Neuroimaging results suggests self-projection and self-reference are managed by a brain network including the medial parietal lobe, the temporoparietal junction (TPJ), and medial frontal cortex (mFC). Interestingly, no significant difference was found between the neural processing of significant and non-significant others. Frontal regions (medial prefrontal and anterior cingulate cortices) showed higher activity for the self POV compared to all other conditions, while all regions showed higher activity for personally closer conditions (self, significant and non-significant others) than the famous-person condition. RSA analysis identified the retrosplenial cortex (RSC) to mediate the representation of cognitive distances for each POVs social network.

As humans we can “self-project” ourselves to different times and locations. The cognitive capacity of self-projection has also been described in the social domain as postulated by the ‘simulation theory’ and ‘theory of mind’ (ToM) (Buckner & Carroll, 2007; Schurz, Kogler, Scherndl, Kronbichler, & Kühberger, 2015; Waytz & Mitchell, 2011). These theories involve (1) the acknowledgment that other people experience their own inner world (unlike toddlers or people on the autistic spectrum) (Frith & Frith, 2003; Leslie, 1987; Premack & Woodruff, 1978), and (2) that understanding that one is capable of putting herself “in the shoes” of another person in order to reason about her mental world including mental states, beliefs or emotions (Gordon, 1986; Harris, 1992). Notably, to mentally adopt the POV of another person one has to use her own self-experience for this situation as a fundamental reference (Tamir & Mitchell, 2010; Waytz & Mitchell, 2011). On top of this, self-projection has been suggested to require a shift of the reference frame, which may be mapped in a metrical-like manner. This hypothesis has been supported by studies demonstrating the similarity between the underlying brain networks in the domains of temporal and spatial cognition (Shahar Arzy, Collette, et al., 2009; Gauthier et al., 2018; Gauthier & van Wassenhove, 2016b; see also a meta-analysis by Spreng, Mar, & Kim, 2009), as well as by the current study which focused on the domain of social cognition. To further explore the relations between our results in the social domain and results involving the time domain, we performed a meta-analysis on the peak voxel of our RSA results using the Neurosynth tool (Yarkoni et al., 2011). This analysis revealed this voxel most strongly associates with studies related to autobiographical memory (Table 2). This association may hint on the role of self-projection and mental-travel in autobiographical memory (Buckner & Carroll, 2007; Tulving, 1985) Focusing on works with social aspects, the most highly-associated cognitive faculties included perspective taking (e.g. Bhanji & Beer, 2012; Jankowski et al., 2014), and moral judgment (Harenski, Antonenko, Shane, & Kiehl, 2008; Hayashi et al., 2014), which may intimately involve the ability to self-project oneself to the POV of the other involved to account for the ethical situation in question (“That which is hateful to you do not do to another”). In this vein, further research on the juxtaposition between memory and self on the social domains was highlighted as an important forefront in neurocognition (Amodio, 2019; Prebble, Addis, & Tippett, 2013). Notably, the region identified in the RSA is adjacent to visual areas, involved also in visual imagery (Ishai, Haxby, & Ungerleider, 2002; Lambert, Sampaio, Scheiber, & Mauss, 2002). The identified region may therefore relate to the appearance of the people imagined rather than self-projection, which should be regressed out in future studies.

**Table 2.** Neurosynth-based meta-analysis. The top ten content-related topics associated with voxels at the left precuneus found in our searchlight-RSA results are shown, as indicated by a Neurosynth based meta-analysis. Topics describing the anatomical regions themselves (e.g. posterior cingulate, retrosplenial, medial, retrosplenial cortex) are not mentioned. Note that 9/10 of these topics are related to memory functions.

| Name | Individual voxel |  | Seed-based network |  |
| --- | --- | --- | --- | --- |
|  | z-score | Posterior prob. | Func. conn. (r) | Meta-analytic coact. (r) |
| autobiographical | 10.7 | 0.88 | 0.49 | 0.52 |
| episodic | 9.69 | 0.81 | 0.44 | 0.49 |
| episodic memory | 8.49 | 0.81 | 0.37 | 0.42 |
| semantic memory | 8.12 | 0.86 | 0.18 | 0.24 |
| retrieval | 7.22 | 0.74 | 0.32 | 0.36 |
| memory | 6.85 | 0.69 | 0.2 | 0.29 |
| memory retrieval | 6.73 | 0.8 | 0.28 | 0.32 |
| concrete | 6.58 | 0.85 | 0.06 | 0.13 |
| autobiographical memory | 6.02 | 0.85 | 0.41 | 0.44 |
| recollection | 5.7 | 0.81 | 0.23 | 0.26 |

‘Self-reference’ is usually defined as the enhanced performance observed for self-related stimuli (Cunningham, Turk, Macdonald, & Neil Macrae, 2008; Leshikar, Dulas, & Duarte, 2015; Rogers, Kuiper, & Kirker, 1977; Symons & Johnson, 1997), since the “self” may serve as a well-organized and frequently used construct (Symons & Johnson, 1997). In young adults, self-reference effect was found for future events with respect to past ones. Namely, participants responded faster to future events compared to past events, perhaps reflecting a future-oriented thinking mechanism (Arzy et al., 2008, 2009). Interestingly, this effect is reversed in older adults, and may also be manipulated by deviations in spatial attention and writing direction (Anelli, Ciaramelli, Arzy, & Frassinetti, 2016b; Anelli et al., 2018). Future studies may investigate whether self-reference in the person domain as found here is sensitive to such manipulations as well.

With respect to functional neuroanatomy, brain activity in the self, significant and non-significant others POV conditions (personally related) differed from the famous-person POV condition (personally unrelated) in all involved clusters. This may be related to the more concrete aspects of the personally related people, as is also postulated by the construal-level theory (Trope & Liberman, 2010). According to this theory social distance may be considered as an egocentric psychological distance that characterizes the level of abstraction of a mental construct (Liviatan, Trope, & Liberman, 2008; Trope & Liberman, 2010). Another remarkable neuroanatomical finding involved the similarity in neural activity for significant and non-significant others. Since, significant others take a major role in one’s mental life, it has been hypothesized that processing significant others require self-related neurocognitive mechanisms unlike non-significant others. Previous studies found stronger activation in the retrosplenial cortex (as found in our RSA) when participants referred to more significant others, suggesting the existence of specific “schemata” (organized self-related models) for the former (Brashears, 2013; Wlodarski & Dunbar, 2016). In the current study, behavioral results showed similar patterns for both self and significant others, possibly due to the intimate acquaintance of subjects with the internal world of the latter. In the neuroanatomical level however, significant and non-significant others showed similar patterns of activations. This finding may explain why neurological disorders regarding significant others are so rare. We speculate that despite their psychological importance, they do not rely on self-related neurocognitive processes and are therefore somehow “immune” to self-related disorders.

The current study aimed to increase ecological validity by considering each participants’ social network, as well as the social networks of participants’ significant- and non-significant-other. Consequently, this resulted a tradeoff with well-controlled homogeneous comparisons between participants. Moreover, three participants chose their parent as their significant other, while six chose their partner and six a close-friend. While comparing between these three categories may be of importance, the objective of the current study was to focus on the performance of self-projection in between the four experimental conditions. Fine-graining of these main conditions may be investigated in future studies. With respect to our selection of Israeli PM Benjamin Netanyahu to act as the famous-person with no personal ties, a stranger may have been a better choice for the study design, yet our ecologically valid task required acquaintance with the persons’ social network. The acquaintance of our participants with Netanyahu may vary, partly due to different political ideology and engagement or sympathy to this specific person. Nevertheless, this was controlled using a designated questionnaire and analyzed in the framework of the RSA. Study participants were relatively young university students, possibly affecting the self-projection effect, as significant-other relationships tend to mature with age (Luong, Charles, & Fingerman, 2011). Participants age may have also influenced the non-significant other chosen, since friendships change with time as well. Nonetheless, setting this age exclusion criteria enabled us to keep homogeneity in between study participants.

In conclusion, our results suggest effects of self-projection and self-reference in the social (person) domain, as demonstrated in previous research dedicated to the space and time domains. These findings may suggest not only a similar neurocognitive mechanism to underlie the three domains but also mutual effects in between the domains of space, time and person that underlie our experiential life and merit further investigations.

## Acknowledgments

The study was supported by the Israel Science Foundation (**Grant No. 1306/18**). We thank Amnon Dafni-Merom for important comments on the manuscript.

## Supplementary materials

**Figure S 1.**
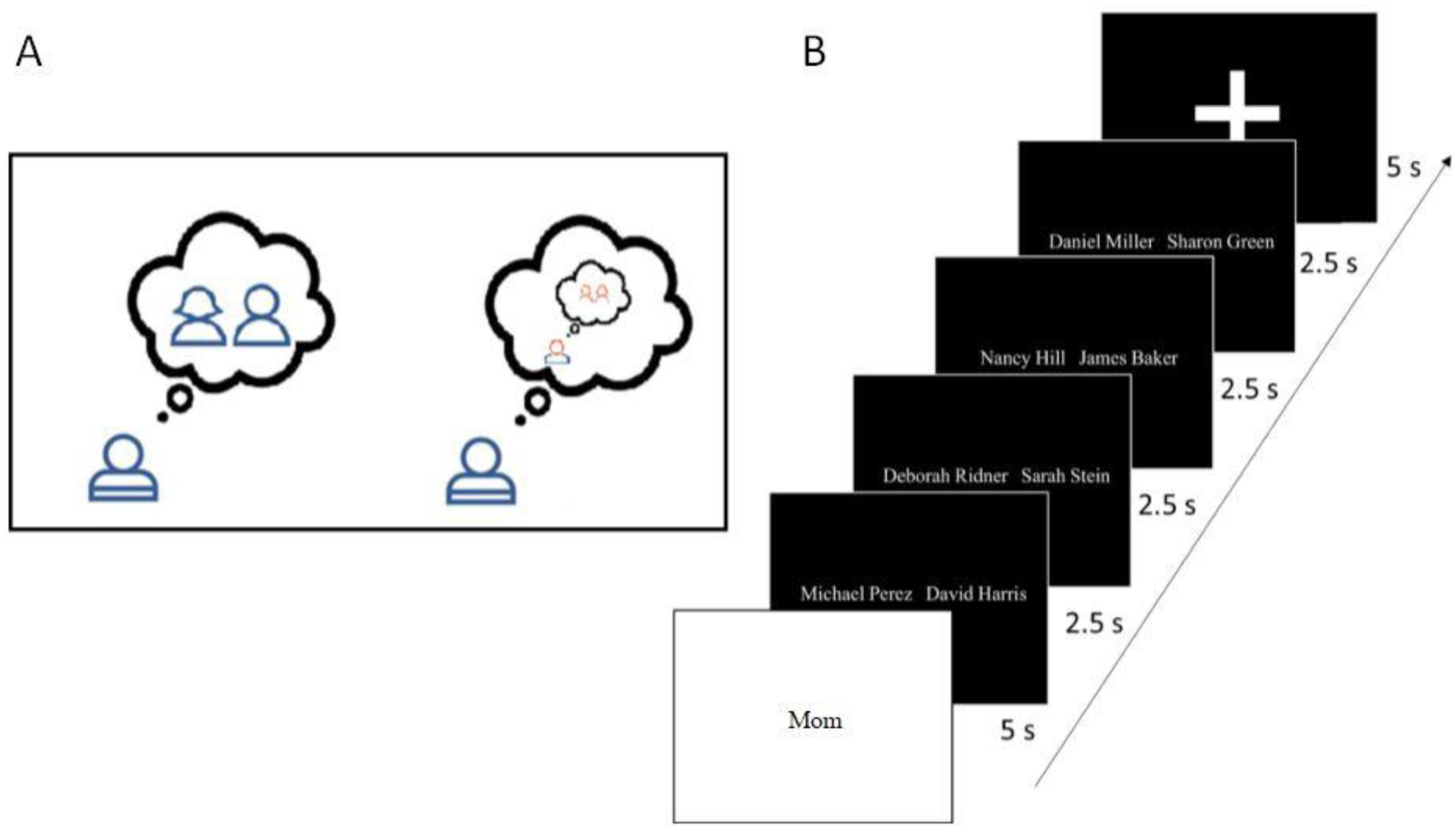
Study Paradigm. (A) Participants were presented with two stimuli, depicting two people from their own (left) or other (significant other, non-significant other, famous-person), and were asked to indicate which of these two people is socialy closer to the imagined POV. (B) Stimuli pairs were presented in a randomized block design. Each block started by presentation of a target stimulus, followed by consecutive presentation of four stimuli pairs. To clean the data from mistyped responses, trials with a response time of less than 300 milli-second (31 responses, approximately 1% of all responses) were excluded, correspondingly to previous study showed high mental processing is not reasonable in such short time (Jain, Bansal, Kumar, & Singh, 2015).

**Figure S 2.**
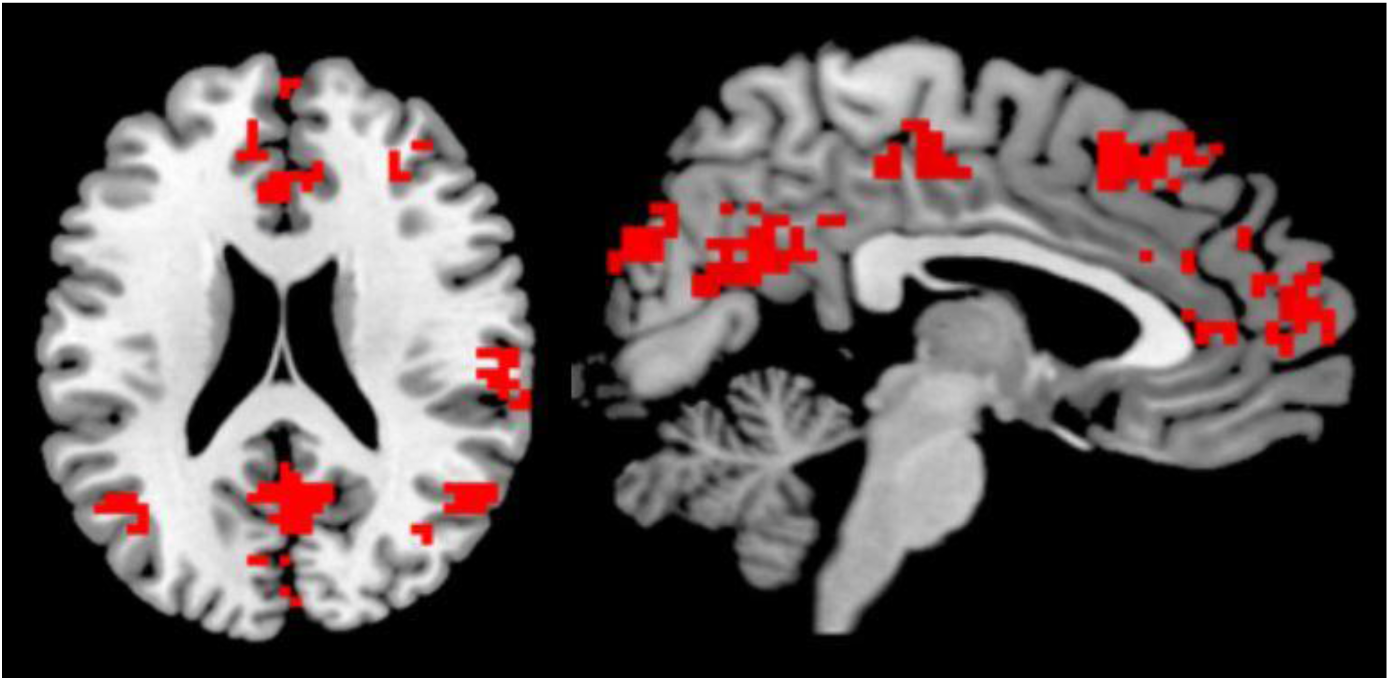
MVPA on self-reference effect. Result of searchlight MVPA analysis with SVM classifier distinguishing between “close” and “distant” stimuli (P<0.001, with cluster size threshold of 20 voxels). activations at the medial prefrontal cortex, anterior and middle cingulate cortex, bilateral temporo-parietal junctions, precuneus, cuneus and bilateral middle frontal cortices show to be sensitive to self-reference effect (relative-distance of stimuli from the adopted POV). A “close” pair consisted from stimuli rated between 1-3 from the relevant POV; in a “distant” pair both stimuli were rated between 4-6.

**Figure S 3.**
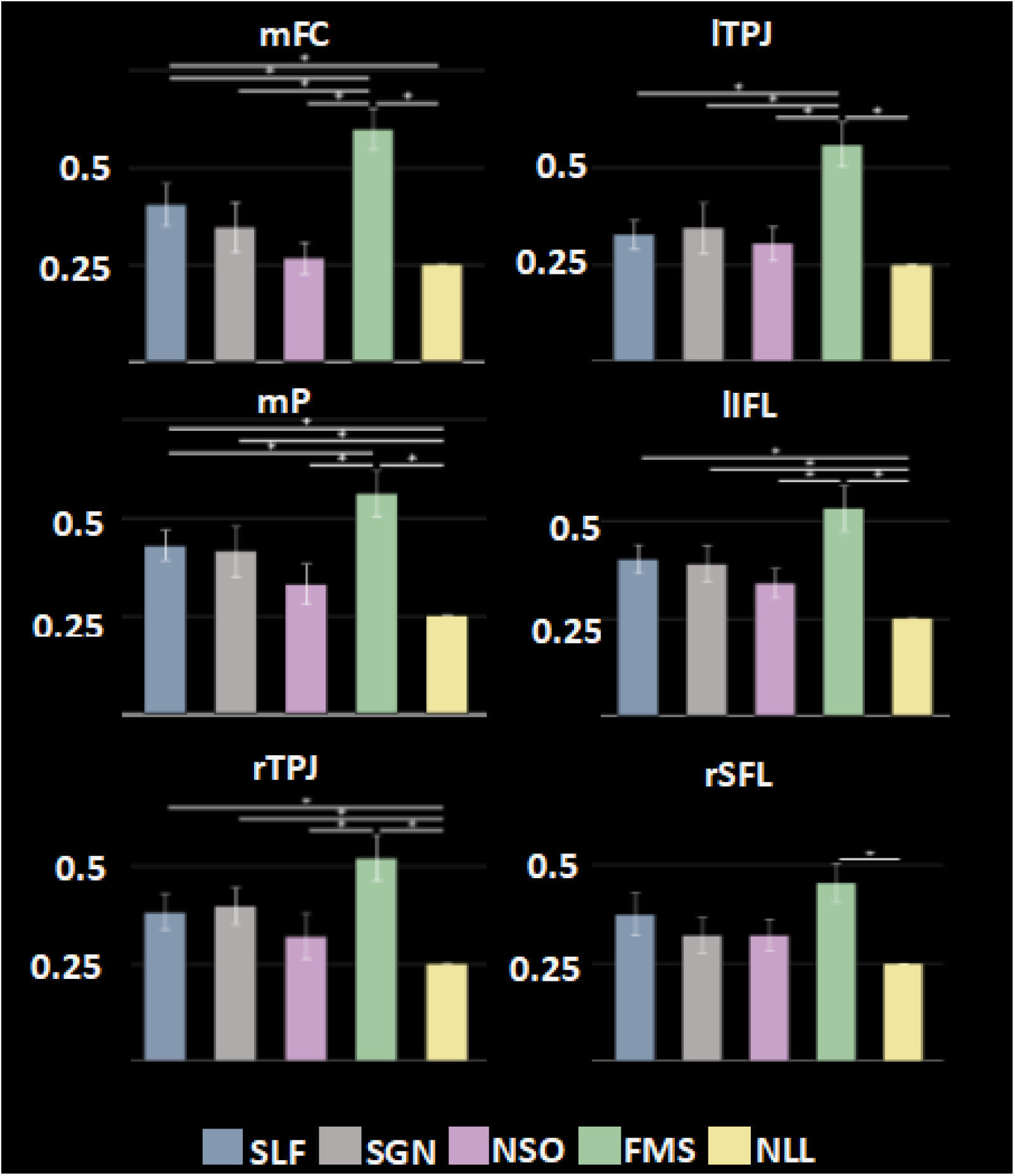
Confusion matrices analysis: F1 scores (F1 considers both the positive predictive value and the sensitivity of the applied MVPA classification) are shown in each ROI, as compared also to a null score (0.25). Significant differences are indicated by asterisks (p<0.05, FWER corrected; mFC – medial frontal cortex, mP – medial parietal, TPJ – temporoparietal junction, IFL – inferior frontal lobe, SFL – superior frontal lobe, r – right, l- left).

**Table S1.**
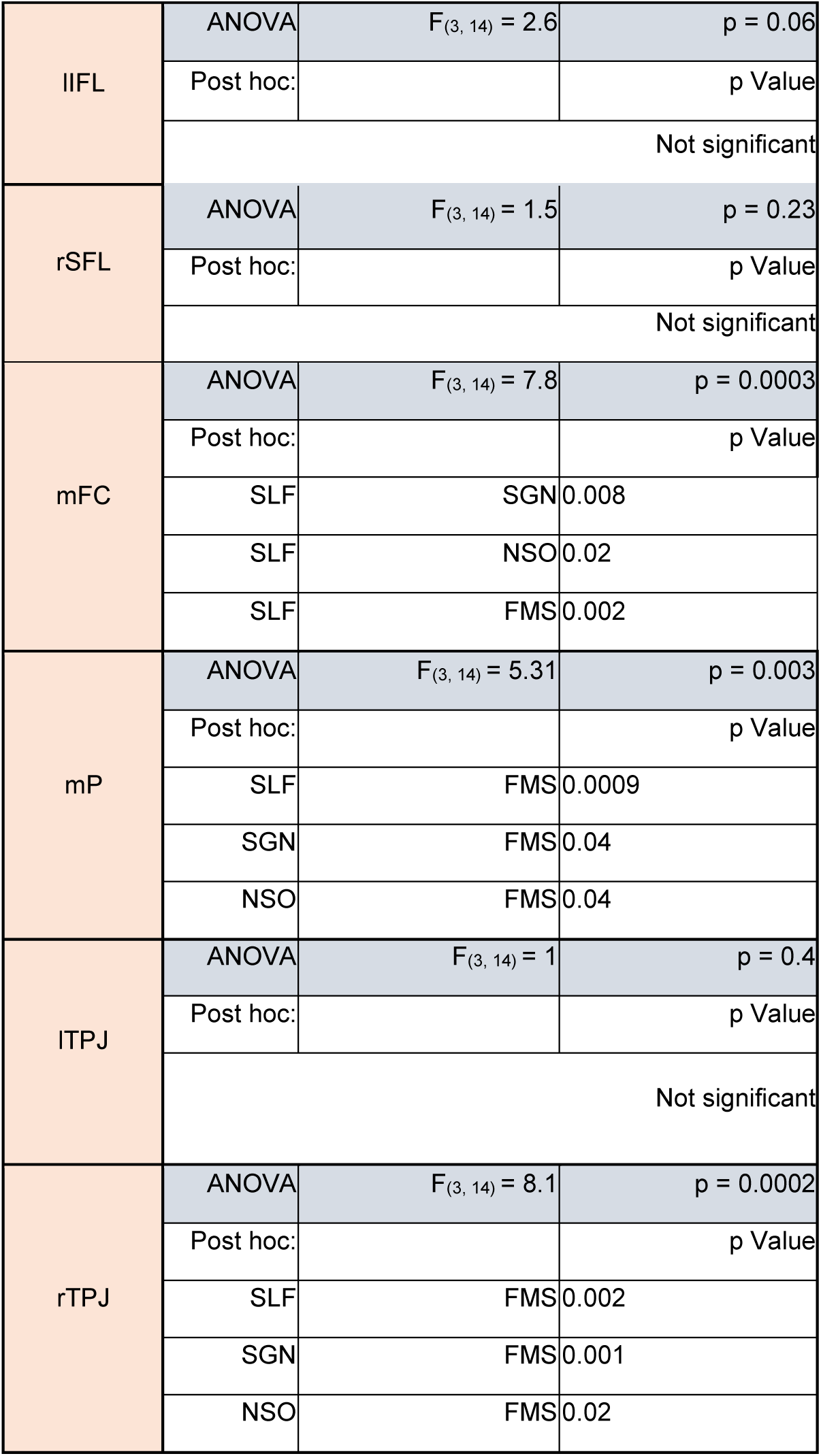
ANOVA and post hoc results for the GLM parameters in each ROI. (mFC – medial frontal cortex, mP – medial parietal, TPJ – temporoparietal junction, IFL – inferior frontal lobe, SFL – superior frontal lobe, r – right, l- left).

**Table S2.** Mean classifier confusion matrices in all ROIs. Each number depicts classifications distribution for a single POV in a given ROI. The diagonal depicts the percentage of right “guesses”. Significant guesses (p < 0.05, Bonferroni corrected) are shown in bold. (mFC – medial frontal cortex, mP – medial parietal, TPJ – temporoparietal junction, IFL – inferior frontal lobe, SFL – superior frontal lobe, r – right, l- left).

| ROI | POV | Classified as: |  |  |  |
| --- | --- | --- | --- | --- | --- |
|  |  | SLF | SGN | NSO | FMS |
| rSFL | SLF | **44%** | 20% | 20% | 16% |
|  | SGN | 32% | 30% | 25% | 14% |
|  | NSO | 31% | 20% | 31% | 17% |
|  | FMS | 22% | 19% | 15% | **44%** |
| lIFL | SLF | **45%** | 16% | 25% | 14% |
|  | SGN | 21% | 37% | 28% | 14% |
|  | NSO | 28% | 26% | 34% | 12% |
|  | FMS | 28% | 9% | 12% | **51%** |
| mFC | SLF | 41% | 21% | 27% | 10% |
|  | SGN | 20% | 36% | 32% | 12% |
|  | NSO | 25% | 35% | 28% | 12% |
|  | FMS | 12% | 16% | 14% | **59%** |
| mP | SLF | **49%** | 20% | 25% | 6% |
|  | SGN | 25% | 40% | 23% | 12% |
|  | NSO | 37% | 15% | 32% | 16% |
|  | FMS | 20% | 12% | 13% | **55%** |
| lTPJ | SLF | 35% | 30% | 25% | 10% |
|  | SGN | 28% | 36% | 22% | 14% |
|  | NSO | 29% | 27% | 28% | 16% |
|  | FMS | 14% | 14% | 15% | **56%** |
| rTPJ | SLF | **40%** | 21% | 26% | 12% |
|  | SGN | 24% | **40%** | 22% | 14% |
|  | NSO | 32% | 18% | 32% | 17% |
|  | FMS | 14% | 22% | 14% | **50%** |

